# Real-time tracking of pupil-phase fluctuations reveals state-dependent modulation of temporal attentional capacity

**DOI:** 10.64898/2026.05.17.725605

**Authors:** Yuta Suzuki, Hsin-I Liao

**Author notes:** Corresponding author: Yuta Suzuki.

## Abstract

Pupil diameter is widely used as an index of arousal and brain state, yet it remains unclear whether slow pupil-linked state fluctuations systematically modulate the effective capacity of temporal attention. Here, we tested this question using an auditory attentional blink paradigm, in which participants were required to detect the first (T1) and second (T2) targets. In Experiment 1, trial-by-trial analyses revealed that successful T2 detection, given correct T1 detection (T2|T1), was associated with smaller baseline pupil size. Furthermore, analyses focusing on slow pupil fluctuations (< 0.2 Hz) revealed that the T2|T1 detection accuracy increased during the pupil dilation phase occurring 0–2 seconds after pupil constriction. In Experiment 2, we used real-time pupillometry to trigger stimulus presentation during predefined phases of ongoing slow pupil dynamics. This closed-loop manipulation produced reliable phase-dependent differences in T2|T1 detection accuracy. Critically, the effect of pupil phase remained significant in a linear mixed-effects model that included baseline pupil size as a covariate, indicating that it cannot be explained by baseline pupil size alone. Together, these findings demonstrate that temporal attentional capacity is shaped not only by arousal level but also by the phase of slow pupil-linked brain-state fluctuations. Our results suggest that the attentional blink reflects a dynamically regulated, state-dependent limitation rather than a fixed processing bottleneck.

## Introduction

Pupillometry has emerged as a powerful tool for tracking moment-to-moment fluctuations in cognitive state. Pupil diameter is widely regarded as an indirect index of arousal level and has been linked to activity in the locus coeruleus–noradrenergic (LC–NA) system (Aston-Jones and Cohen, 2005; Murphy et al., 2014; Reimer et al., 2016).

Several studies have reported negative correlations between changes in pupil diameter (i.e., baseline pupil size) and performance in perceptual and cognitive tasks, such that smaller baseline pupil size is associated with better behavioral outcomes or perceptual stability (Orden et al., 2000; Kristjansson et al., 2009; Grandchamp et al., 2014; Mittner et al., 2014; Hopstaken et al., 2015; Suzuki et al., 2022). For example, van Kempen and colleagues reported that smaller baseline pupil sizes were linked to faster reaction times and higher accuracy in a moving-dot detection task (Kempen et al., 2019). Together, these findings suggest that the brain state reflected in baseline pupil size is systematically related to cognitive performance across domains.

The attentional blink (AB) provides a particularly sensitive indicator of fluctuations in effective attentional capacity. When two targets are presented in close temporal proximity within a rapid stream, identification of the second target (T2) is often impaired if it follows the first target (T1) within tens to hundreds of milliseconds (Raymond et al., 1992). Contemporary accounts attribute the AB to a temporary bottleneck in post-perceptual consolidation processes, whereby T1 occupies a limited-capacity stage required for reportable encoding (Chun and Potter, 1995; Wyble et al., 2009). Importantly, AB performance varies substantially across trials even under identical stimulus conditions (Martens and Wyble, 2010). This variability suggests that the efficiency of consolidation is not fixed but instead may depend on fluctuations in underlying brain state.

Importantly, attentional capacity fluctuates over time as a function of ongoing brain state. Rather than reflecting a static resource limitation, trial-to-trial variability in performance may arise from slow modulations in neural gain and cortical excitability mediated by neuromodulatory systems such as the LC–NA system (Aston-Jones and Cohen, 2005; Sara, 2009). From this perspective, moment-to-moment changes in internal state dynamically alter the effective availability of attentional resources, thereby shaping perceptual competition and consolidation processes. Slower fluctuations in pupil size index dynamic changes in brain state over seconds (Joshi et al., 2016; Gee et al., 2017). These slow pupil-linked fluctuations (< 0.2 Hz) are coupled with large-scale cortical dynamics, including infra-slow BOLD fluctuations and modulations of alpha-band power envelope (Yellin et al., 2015; Montefusco-Siegmund et al., 2022). Notably, pupil size correlates negatively with activity in sensory cortices and positively with activity in default mode network regions (Yellin et al., 2015), suggesting that pupil-linked brain state reflects structured reconfigurations of large-scale brain networks rather than global activation. Furthermore, recent work indicates that stimulus representations themselves vary as a function of the phase of pupil-linked brain-state fluctuations (Yang et al., 2026), underscoring the functional relevance of these dynamics. However, it remains unclear whether these slow brain-state fluctuations translate into systematic changes in the effective capacity of temporal attention. While several studies have reported baseline pupil size as an index of arousal level related to behavioral performance, it remains unclear whether the probability of successfully encoding temporally competing stimuli depends on the phase of ongoing slow pupil-linked brain state dynamics.

In the present study, we employed an auditory attentional blink paradigm (Shen and Mondor, 2006) to minimize confounding effects of visual stimulation on pupil change and to isolate central, modality-independent attentional limitations. Although the attentional blink was first reported in the visual domain, subsequent studies demonstrated its presence in the auditory domain, suggesting that it reflects a modality-independent limitation in temporal attentional capacity (Duncan et al., 1997; Shen and Mondor, 2006). In Experiment 1, we examined whether baseline pupil size and the phase of slow pupil fluctuations below 0.2 Hz during the stimulus presentation predicted target detection. Because these brain state related dynamics unfold continuously over several seconds, understanding their influence on temporal attention requires tracking pupil fluctuations in real time and relating behavioral performance to specific phases of ongoing state transitions. In Experiment 2, we monitored pupil size in real time to trigger stimulus presentation during either the dilation or constriction phase of ongoing pupil fluctuations, thereby testing whether specific phases of slow pupil dynamics provide an optimal state for target detection. By distinguishing between level-dependent effects of pupil-linked arousal (baseline pupil size) and dynamics-dependent effects (phase-specific slow fluctuations), the present study aimed to clarify how slow brain-state dynamics modulate temporal attention and consolidation efficiency in the attentional blink.

## Method

### Participants

A total of 51 participants (32 men and 19 women, aged 20–40 years, mean age = 25.31 years, SD = 5.48), with normal or corrected-to-normal vision, took part in the experiments. Twenty volunteers participated in Experiment 1. Two participants were excluded as outliers since their T1 and T2 detection accuracy or false alarm (FA) rate exceeded a robust outlier criterion. Specifically, we first computed the median of T1, T2 detection accuracy and FA across participants. When the median was 1.0 (i.e., ceiling effects), we applied a z-score criterion based on the mean and standard deviation and excluded participants with ∣ ∣>3. Otherwise, we used the modified z-score based on the median absolute deviation (MAD) and excluded participants with ∣ *M*z ∣>3.5 (Iglewicz & Hoaglin, 1993). Thirty-one volunteers participated in Experiment 2. Five participants were excluded using the same criterion described above. All experimental procedures were approved by the Research Ethics Committee of Nippon Telegraph and Telephone (NTT) Communication Science Laboratories (R5-001) and were conducted in accordance with its ethical guidelines. All participants provided written informed consent.

### Auditory stimulus

The auditory attentional blink (AAB) paradigm was adapted from Shen and Mondor, (2006). All pure tones were 30 ms in duration and included 2-ms linear onset and offset amplitude ramps to eliminate clicking artifacts. Twenty-one pure tones were used as distractors. The frequencies of these tones were logarithmically spaced and ranged from 529 to 1330 Hz. The specific frequencies were 529, 554, 580, 607, 636, 666, 697, 730, 764, 800, 838, 877, 918, 961, 1006, 1056, 1106, 1158, 1213, 1270, and 1330 Hz. The target tone (T1) consisted of six 5-ms pulses and could take any of the 21 frequencies used for the distractor tones. The probe tone (T2) was a frequency glide that smoothly changed from 636 to 1006 Hz. Each tone was presented for 30 ms followed by a 50 ms silent interval. Using these tones, four types of sequences were used in the experiments: a non-target sequence, a T1-only sequence, a T2-only sequence, and a T1T2 sequence (see Fig. 1A). In the T1- and T2-only sequences, the fourth or sixth sound in the sequence was replaced with the T1 or the T2, respectively. In the T1T2 sequence, both T1 and T2 were presented within the same sequence. Within the T1T2 sequence, five lag conditions (Lag 1–Lag 5) were defined based on the temporal distance between T1 and T2. In Lag 1, T2 was presented immediately after T1. In Lag 5, T2 was presented after four intervening distractor tones following T1. Each sequence consisted of 16 tones. The T1-only, T2-only, and T1T2 sequences were always followed by a non-target sequence, resulting in a total sequence duration of 2560 ms (i.e., (30 + 50) x 32 = 2560).

**Figure 1.**
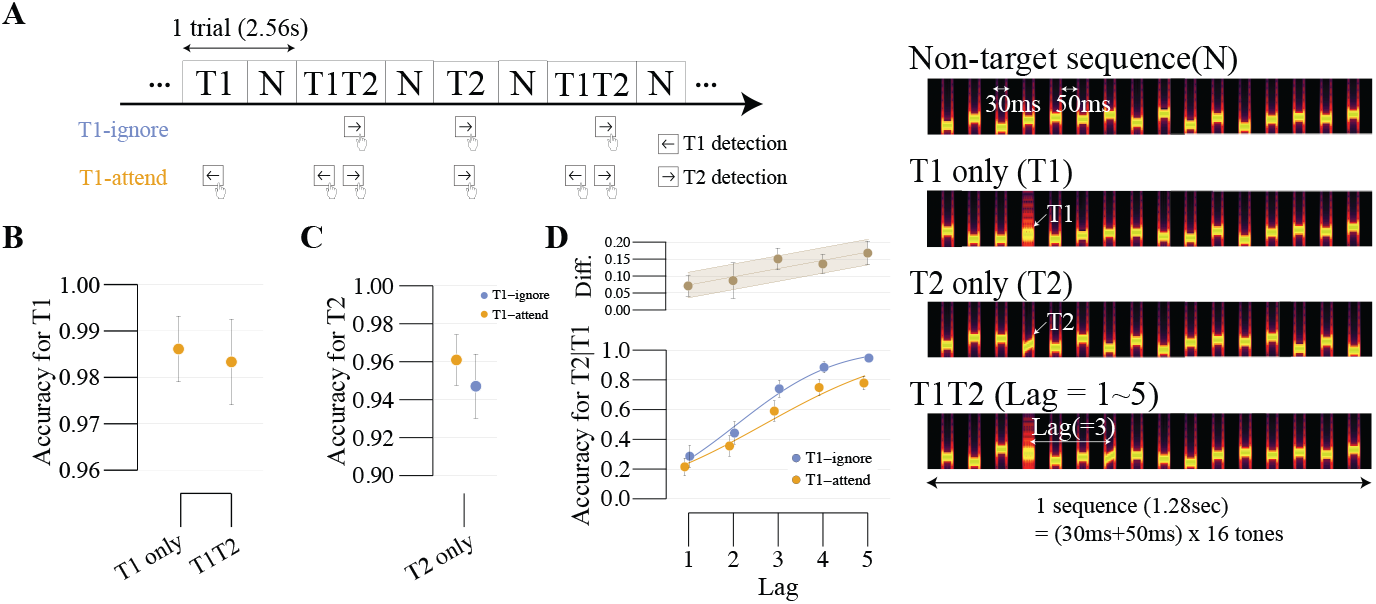
Schematic of the auditory attentional blink task and behavioral performance. (A) Schematic illustration of the task. Each trial consisted of a 2.56-s auditory sequence containing non-target sounds (N-sequence), only T1 sound (T1-sequence), only T2 sound (T2-sequence), or both target sounds (T1T2-sequence). In the T1-ignore condition, participants detected only T2 sound, whereas in the T1-attend condition they detected both T1 and T2 sound. Example spectrograms of the N-, T1-, T2-, and T1T2-sequences are shown on the right. **(B)** T1 detection accuracy in the T1- and T1T2-sequence. **(C)** T2 detection accuracy in the T2-only sequence. **(D)** Lag-dependent performance in the T1T2-sequence. The lower panel shows T2 detection accuracy in the T1-ignore (blue) and the T1-attend (orange) condition. The upper panel shows the performance difference between the two conditions. Error bars and shaded areas represent the

### Experimental apparatus

The stimulus was generated digitally with a sampling rate of 44.1 kHz and presented diotically via headphones (MDR-7506, Sony, Tokyo, Japan) at a sound pressure level of 65 dB. The luminance of the screen was measured by a spectroradiometer (CS-3000HDR, KONICA MINOLTA, Tokyo, Japan). An LCD monitor (Flex Scan EV2460-BK, EIZO, Ishikawa, Japan) with a resolution of 1920 × 1080 and a refresh rate of 60 Hz was used for eye-tracker calibration. During the experiment, the background screen was always black (0.025 cd/m^2^) with CIE xy coordinates of (0.315, 0.333). The fixation cross was located at 0.1 degrees from the center (29.95 cd/m^2^) with a line width of 1 pixel. The task was conducted in a bright room (30.21 lx) and executed implemented in Python 3.10 using PsychoPy (Peirce, 2007).

### Experiment 1

#### Procedure

Each participant’s head was stabilized using a chin rest to minimize head movements at a viewing distance of 600 mm from the computer screen. At the beginning of each block, a standard five-point calibration provided by the EyeLink software (v5.15) was performed. Before the main experiment, participants completed a training session to familiarize themselves with the auditory targets T1 and T2. The experiment was divided into four sessions in total. Two task conditions of T1-attend and T1-ignore were included. In the T1-attend condition, participants were required to report both T1 and T2 when detected. In the T1-ignore condition, participants were instructed to ignore T1 and respond only to T2. Participants responded using a keyboard, pressing the left arrow key for T1 and the right arrow key for T2 when they detected the T1 target and T2 target, respectively. The T1-attend and T1-ignore conditions were each repeated twice, and the order of the sessions was counterbalanced across participants. Each session consisted of 120 auditory sequences (see the “Auditory stimulus” section) which included 20 T1 sequences, 20 T2 sequences, and 80 T1T2 sequences (16 sequences for each lag condition, Lag 1–5). Participants took a short break between sessions.

Participants were instructed to maintain fixation on a central fixation cross throughout the experiment in order to ensure stable pupillary measurements. Since each sequence lasted 2.56 seconds, one session lasted 307.2 seconds (5.12 minutes).

### Pupil recording and preprocessing

Pupil diameter and eye movements during the task were measured using an eye-tracking system (EyeLink 1000, SR Research, Oakland, Canada) at a sampling rate of 1000 Hz. Eye movements from each participant were recorded using an infrared video camera at a resolution of no more than 0.1°. For each participant, data from the eye with the least data loss, either the left or right eye, were used for analysis. The tracking was performed based on pupil diameter using the ellipse mode throughout the study. Pupil diameter was generated by the eye-tracker in arbitrary units and converted to millimeters from pixels. Pupil samples during eye-blinks, which were obtained as the zero values in the data and samples falling outside the 99% confidence interval of the distribution for the entire experiment for each participant were interpolated using cubic Hermite interpolation.

### Pupil phase analysis

After interpolation, a fourth-order low-pass filter with a cutoff frequency of 0.15 Hz was applied to the pupil time series. Following filtering, pupil events (pupil dilation and constriction) were detected from the smoothed signal using a peak–trough detection algorithm; candidate extrema were then identified by examining sign changes in the first temporal derivative of the signal. Specifically, a peak was defined as a point where the derivative changed from positive to negative, whereas a trough was defined as a point where the derivative changed from negative to positive. To suppress spurious detections caused by noise, extrema were accepted only if the magnitude of the pupil change exceeded a predefined threshold parameter of 0.001 pixels. The algorithm identified local extrema corresponding to either pupil troughs or peaks depending on the experimental mode. Two analysis modes were implemented: pupil dilation phase (PD phase) based on the detection of pupil troughs and pupil constriction phase (PC phase) based on the detection of pupil peaks.

### Statistical analysis

The level of statistical significance was set to p < 0.05 for all analyses. Pairwise comparisons of the main effects were corrected through multiple comparisons using the modified Bonferroni method. Effect sizes were reported as partial η^2^ (η^2^_p_) for ANOVA and Cohen’s d (*d*_*z*_) for t tests. To quantify the evidence in the data, we performed Bayesian one-sample t-tests using the BayesFactor package (v0.9.12-4.7) (Morey, 2018) for the R software (Version 4.5.1) (RCoreTeam, 2016). We report Bayes factors as BF_10_ estimating the relative weight of the evidence in favor of H_1_ over H_0_ as BF_10_. Greenhouse-Geisser corrections were performed when the results of Mauchly’s sphericity test were significant.

### Psychometric function fitting and bootstrap

We quantified how conditional probe detection performance changed with temporal lag by fitting a psychometric function to the relationship between lag condition and conditional probe detection probability, P(T2|T1). Data were grouped by phase condition (i.e., the PC phase vs. the PD phase), and for each group we fitted a cumulative normal function with lag condition as the predictor and P(T2|T1) as the dependent variable. We derived the threshold as the lag value at which predicted performance reached a criterion probability of 0.5 (i.e., the 50% point on the fitted curve). Parameter estimation and threshold extraction were conducted using quickpsy package (v0.1.5.1) (Linares and López-Moliner, 2016). To quantify uncertainty in the threshold estimate, we performed a parametric bootstrap with 10,000 resamples (B = 10,000). Non-finite bootstrap estimates were excluded prior to summarization. Model fitting was performed independently for each analysis group.

### Experiment 2

#### Procedure

Calibration and training were conducted before the experiment following the same procedure as in Experiment 1. The experiment consisted of six sessions in total. Throughout the experiment, participants were required to respond whenever they detected either T1 or T2. The PC- and PD-phase conditions were each repeated three times, and the order of the sessions was counterbalanced across participants. One session continued until 60 target sequences had been presented, consisting of 5 T1 sequences, 5 T2 sequences, and 50 T1T2 sequences (10 sequences for each lag condition, Lag 1–5). Each session lasted approximately 7–8 minutes. Participants took a short break between sessions. Pupil diameter was analyzed in real time using a custom Python program according to the following procedure.

### Real-time pupil processing

Incoming pupil samples were stored in a queue structure and processed in a separate analysis thread. Real-time analysis began 60 s after the onset of the first sound stimulus. Pupil samples were accumulated at a sampling rate of 100 Hz and smoothed using a 15-s moving-average sliding window. Raw pupil diameter signals were first subjected to outlier rejection. The first derivative of the pupil signal was computed, and the interquartile range (IQR) of the derivative distribution was used to identify extreme changes. Specifically, the 25th and 75th percentiles of the derivative were calculated, and samples outside 1.5 × IQR were removed to exclude large artifacts such as blinks or tracking errors. These samples were replaced with zeros and subsequently interpolated by cubic interpolation. After interpolation, a fourth-order low-pass filter with a cutoff frequency of 0.15 Hz was applied to the pupil time series. The same pupil phase detection algorithm as in Experiment 1 was then employed. Because extrema were determined based on sign changes in the derivative, confirmation of a peak or trough required several subsequent samples after the true extremum occurred. Consequently, stimulus triggering occurred with an inherent delay of approximately 1 s relative to the actual pupil extremum, corresponding to the timing relationship illustrated in Fig. 4A. This analysis sequence was applied whenever a new pupil sample was detected. When a peak for the PC phase or a trough for the PD phase was detected, a trigger signal was emitted, after which the T1 and/or T2 sounds were presented.

### Regressing out baseline pupil size effects

To estimate the influence of baseline pupil size on target detection performance, we fitted a linear mixed-effects model predicting target detection performance from baseline pupil size, with participant included as a random intercept:

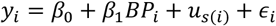

where *BP*_*i*_ denotes baseline pupil size, *u*_*s*(*i*)_ is the subject-specific random intercept, and *ϵ*_*i*_ is the residual error. Residualized performance was then computed as

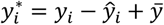

where *ŷ*_*i*_ is the model-predicted value and 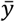 is the grand mean of *y*. Thus, the residualized values 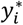 represents behavioral variability after removing the contribution of baseline pupil size. These residualized values were used in subsequent analyses of pupil phase.

## Data and code availability

Participants’ raw data and the Python and R scripts used for data analysis are publicly available at https://github.com/suzuki970/PupilPhase-AttentionalBlink. All data required to evaluate the conclusions of this study are included in the paper and its Supplementary Materials.

## Results

### Experiment 1

#### Behavioral data

First, accuracies for T1 and the probe (T2) were very high, indicating that participants were able to reliably discriminate the target sounds from the distractors (T1: M=0.984, SD=0.039, Probe: M=0.954, SD=0.064; see Figure 1B and 1C). To confirm the attentional blink (AB) effect, we conducted a two-way repeated-measures ANOVA with lag and session (T1-attend vs. T1-ignore) as within-subject factors. The analysis revealed a significant main effect of session (*F*(1,17) = 28.782, p < 0.001, 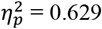 , *BF*_10_ = 4.863). Post hoc analyses showed that, consistent with previous studies, probe detection probability significantly decreased in the T1-attend condition compared to the T1-ignore condition (ps < .05), except at lag 2 (p = 0.115) as shown in Figure 1D. We also observed a significant main effect of lag (*F*(4,68) = 49.99, p < 0.001, 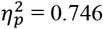 , *BF*_10_ > 100), whereas the interaction between lag and session was not significant (*F*(4,68) = 1.693, *p* = 0.162, 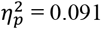 , *BF*_10_= 0.087).

### Relationship between baseline pupil size and T2|T1 detection accuracy

To examine whether baseline pupil size influenced target detection, we compared pupil diameter prior to target onset between hit and miss trials. Pupil time courses around target presentation are shown in Figure 2A. Baseline pupil size was defined as the mean pupil diameter during the 1-second period preceding T2 onset. The baseline pupil size was averaged across lag conditions and separated by response (hit vs. miss), as shown in Figure 2B. The results showed that baseline pupil size was significantly smaller for hit trials compared to miss trials (*F*(1,17) = 6.334, *p* = 0.022, 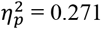 , *BF*_10_ = 0.587). Despite the non-significant interaction (*F*(1,17) = 3.01, *p* = 0.101, 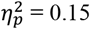 , *BF*_10_ = 0.37), further analysis including Bayes factors revealed that this effect was present only in the T1-attend condition (*t*(17) = 2.911, *p* = 0.01, *d*_*z*_ = 0.146, *BF*_10_ = 5.405), but not in the T1-ignore condition (*t*(17) = 1.269, *p* = 0.222, *d*_*z*_ = 0.052, *BF*_10_ = 0.486).

**Figure 2.**
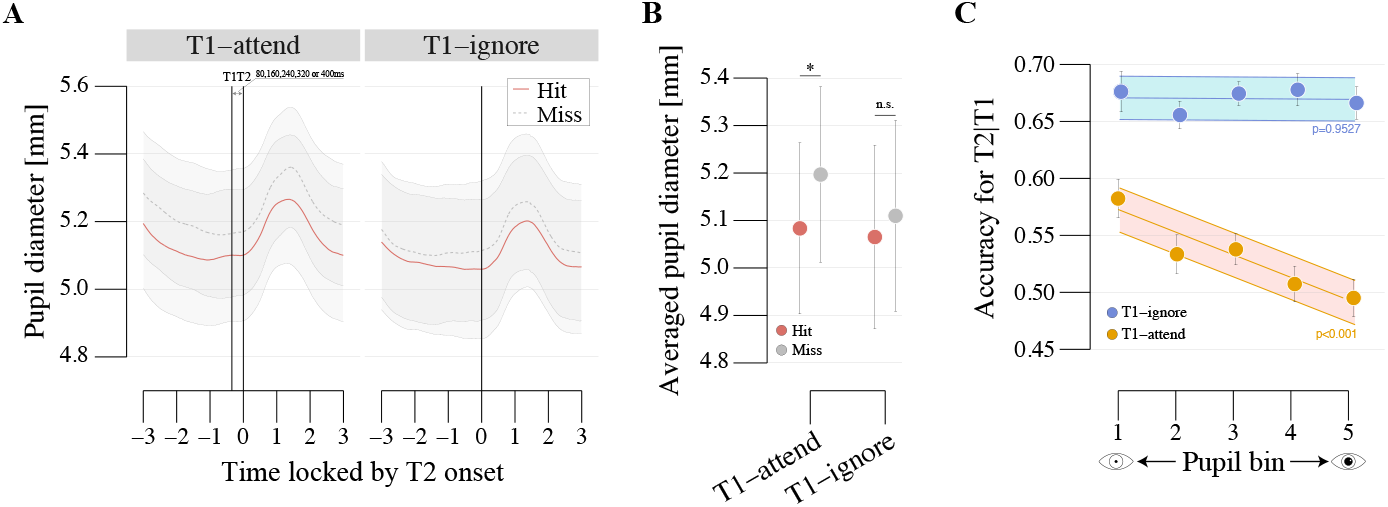
Relationship between baseline pupil size and target detection performance. **(A)** Time course of pupil diameter aligned to T2 onset for hit and miss trials in the T1-attend condition. Shaded areas represent the standard error of the mean across participants. The vertical line indicates the onset of T2. **(B)** Baseline pupil size (1 s before T2 onset) for hit and miss trials in the T1-attend and T1-ignore conditions. **(C)** T2 detection accuracy as a function of baseline pupil size. Trials were sorted according to baseline pupil size and divided into five bins from smallest to largest. T2 detection accuracy is plotted separately for the T1-attend (orange) and T1-ignore (blue) conditions. Error bars and shaded regions represent the standard error of the mean.

Furthermore, to examine the relationship between baseline pupil size and detection performance, trials were sorted according to baseline pupil size and divided into five bins from smallest to largest (following Kempen et al. (2019)). Accuracy for each bin was plotted separately for each attention condition (Figure 2C). We found a significant correlation between baseline pupil size and probe detection probability in the T1-attend condition (p < 0.001), whereas no correlation was observed in the T1-ignore condition (p = 0.9527).

### Relationship between slow pupil change and T2|T1 detection accuracy

Next, we examined low-frequency pupil dynamics by applying a 0.15-Hz low-pass filter to the pupil signal, in line with previous on slow pupil-linked brain-state fluctuations. Figure 3A illustrates an example where pupil constriction (PC) events were indicated by blue markers and Figure 3B (upper panel) shows the PD phase following the PC events. Pupil dynamics following the PC phase is shown in Supplementary Figure 1A. Time interval between each event and T1 onset was calculated. Trials were then grouped into 1-second time bins until 6 seconds relative to these events. We computed the difference between bin-wise T2 detection accuracy for the PD phase collapsed across lags (Figure 3B, lower panel) and separately for each lag (Figure 3C).

**Figure 3.**
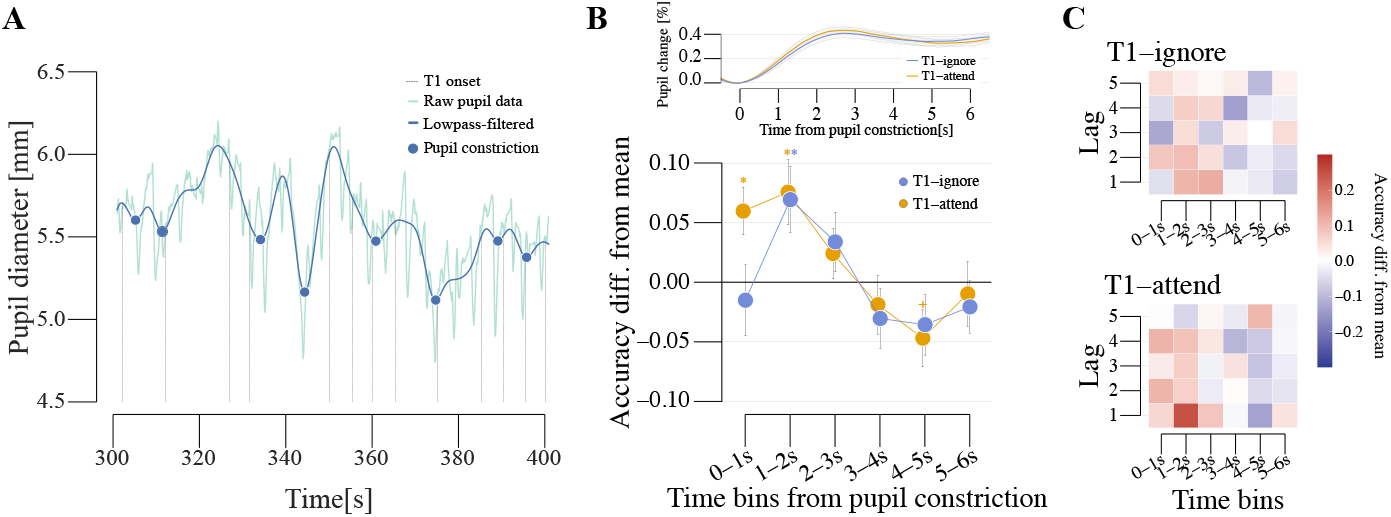
Relationship between phase of pupil dynamics and target detection performance. **(A)** Example time course of pupil diameter during the experiment. The green line shows the raw pupil signal and the blue line indicates the lowpass-filtered signal. Blue markers denote detected pupil constriction events (see “Method”). Vertical black dotted lines indicate the timing of T1 sound onset. **(B)** Modulation of detection accuracy as a function of time relative to pupil constriction (i.e., the PD phase). The upper panel shows the average pupil response aligned to pupil constriction events. The lower panel shows the difference between bin-wise detection accuracy and the overall mean accuracy in the PD phase. Error bars denote the standard error of the mean. Asterisks indicate significant deviations from the mean performance. **(C)** Heatmaps showing lag-dependent changes in detection accuracy in the PD phase for the T1-ignore (top) and T1-attend (bottom) conditions. Each cell represents the deviation in accuracy from the overall mean for a given lag.

**Figure 4.**
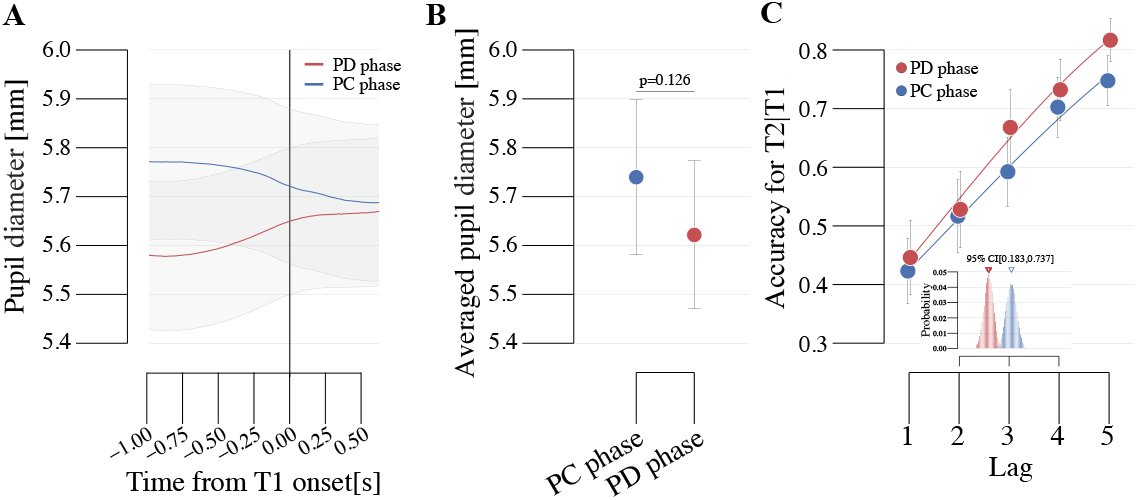
Real-time pupil-phase-triggered stimulation. **(A)** Average pupil diameter around T1 onset for trials triggered during the PD and PC phases. **(B)** Baseline pupil size measured prior to T1 onset for trials triggered during the PD and PC phases. **(C)** T2 detection accuracy as a function of lag for the PD-phase (blue) and PC-phase (red) conditions. The inset shows the bootstrap distribution of the estimated lag thresholds for the two conditions, with the 95% confidence interval of the difference between thresholds indicated. The shaded area and error bars represent the standard error of the mean.

We found that the accuracy for P(T2|T1) significantly improved immediately following the PD phase. Specifically, in the T1-attend condition, accuracy was significantly higher during the 0–1 s (*t*(17) = 2.975, *p* = 0.009, *d*_*z*_ = 0.992, *BF*_10_ = 6.049) and 1–2 s (*t*(17) = 2.757, *p* = 0.013, *d*_*z*_ = 0.919, *BF*_10_ = 4.134). In the T1-ignore condition, a significant improvement was observed in the 1–2 s (*t*(17) = 2.511, *p* = 0.022, *d*_z_ = 0.837, *BF*_10_ = 2.728). In the T1-attend condition, a marginally significant decrease in accuracy was observed during the 4–5 s interval (*t*(17) = -1.968, *p* = 0.066, *d*_*z*_ = -0.656, *BF*_10_ = 1.17). In contrast, no significant changes in accuracy were observed following the PC phase during the 1–2 s interval (*t*(17) = -0.481, *p* = 0.637, *d*_*z*_ = -0.16, *BF*_10_ = 0.269 for the T1-attend; *t*(17) = 1.615, *p* = 0.125, *d*_*z*_ = 0.538, *BF*_10_ = 0.725 for the T1-ignore).

### Experiment 2

#### Behavioral data

To validate the improvement in AB detection following the PD phase observed in Experiment 1, we implemented real-time pupil monitoring and presented T1 at specific phases of slow pupil fluctuations (0.15 Hz), corresponding to the PD or PC phase. Figure 4A shows pupil dynamics immediately preceding T1 presentation in both conditions. As intended, T1 was successfully presented within 0–1 s after the onset of the PD or PC phase. Detection accuracy for the T1- and T2-only sequences remained high (T1: M=0.947, SD=0.097, Probe: M=0.818, SD=0.238; see Supplementary Figures 2A and 2B). We confirmed that mean pupil size around T1 onset did not significantly differ between the PD and PC phases as shown in Figure 4B (*t*(25) = -1.582, *p* = 0.126, *d*_*z*_ = -0.149, *BF*_10_ = 0.622).

### Smaller baseline pupil size was associated with higher T2|T1 detection accuracy

To examine whether the relationship between baseline pupil size and detection performance observed in Experiment 1 could be replicated under real-time pupil-triggered presentation, we repeated the same bin-based analysis in Experiment 2 (see Supplementary Figure 2C). Trials were sorted according to baseline pupil size and divided into five bins from smallest to largest, following the same procedure as in Experiment 1. Detection accuracy for P(T2|T1) was then calculated separately for each bin and phase condition. Consistent with the results of Experiment 1, we again observed a significant correlation between baseline pupil size and probe detection probability in the PC (p < 0.001) phase and PD phase (p = 0.0064) as Supplementary Figure 2C. These results indicate that the relationship between ongoing pupil state and attentional blink performance was reproducible even when stimulus presentation was controlled in real time based on slow pupil dynamics.

### PD-phase triggering improved T2|T1 detection relative to PC-phase triggering

A two-way repeated-measures ANOVA examining the main effect of phase condition revealed that accuracy in the PD phase was significantly higher than in the PC phase (*F*(1,25) = 6.238, *p* = 0.019, 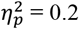 , *BF*_10_ = 0.744), consistent with our prediction based on the phase-triggering manipulation. We estimated the lag threshold (50% of accuracy, corresponding to the conventional midpoint of the psychometric function (Wichmann and Hill, 2001)) from the fitted psychometric functions for each phase condition and assessed its uncertainty via parametric bootstrap (10,000 iterations). Figure 4C shows lag-dependent P(T2|T1) performance. The inset in Figure 4C shows the bootstrap distributions of the threshold estimates for each condition. The 95% bootstrap confidence interval for the threshold difference between the PC and PD phases (PC – PD) was [0.18, 0.735], indicating that the threshold was significantly higher in the PC phase than in the PD phase.

As described in the preceding section, baseline pupil size was significantly associated with detection performance in Experiment 2. We therefore tested whether the improved detection performance during the PD phase could be explained merely by differences in baseline pupil size. To rule out this possibility, we fitted a linear mixed-effects model predicting detection accuracy from baseline pupil size (see “Regressing out baseline pupil size effects”). After removing the contribution of baseline pupil size, the main effect of phase remained significant when residualized detection accuracy was used (*F*(1,25) = 4.625, *p* = 0.041, 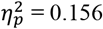 , *BF*_10_ = 0.794). This result indicates that the increased T2|T1 accuracy during the PD phase cannot be explained solely by baseline pupil size.

## Discussion

The present study investigated the relationship between state fluctuations in pupil–brain coupling and temporal attentional capacity. Using an auditory attentional blink paradigm that minimized light-driven changes in pupil diameter, we show that variability in T2 detection accuracy is explained not only by tonic arousal levels, as indexed by baseline pupil size, but also by the phase of slow pupil fluctuations. In Experiment 1, we replicated the classical auditory attentional blink effect, in which T2 detection decreases when T1 and T2 must both be processed. In addition, we found that a smaller baseline pupil size was associated with improved T2 detection accuracy. This finding is consistent with the view that baseline pupil size reflects underlying brain states that constrain post-perceptual integration processes. Moreover, T2 detection probability increased during the PD phase, which occurred several seconds after pupil constriction. In Experiment 2, we leveraged this observation by presenting targets in real time based on detected pupil dynamics. Targets presented during the PD phase were detected more accurately than those presented during the PC phase, demonstrating the feasibility of using real-time pupil monitoring to probe the relationship between slow pupil fluctuations and cognitive state. Collectively, these findings suggest that the capacity limits revealed by the attentional blink are dynamically regulated by ongoing fluctuations in arousal and brain state unfolding over several seconds.

The relationship between tonic arousal dynamics reflected in baseline pupil size and task performance is consistent with previous work linking pupil diameter to activity in the LC– NA system and behavioral performance (Reimer et al., 2014; McGinley et al., 2015). Activation of LC neurons and the resulting release of noradrenaline are known to reorganize neural network dynamics and regulate cognitive performance (Bouret and Sara, 2005). In line with the Yerkes–Dodson law, task performance typically follows an inverted-U relationship with arousal (McGinley et al., 2015). In the present task, larger baseline pupil size was associated with increased attentional blink failures (missed T2 targets), suggesting that relatively smaller baseline pupil size corresponded to a more favorable arousal level for perceptual integration. Notably, the relationship between baseline pupil size and T2 detection accuracy emerged specifically in conditions requiring the detection of both T1 and T2, suggesting that this arousal–performance relationship becomes evident under conditions of increased task demand. Importantly, this pattern was reproduced when stimulus timing was modulated in real time according to pupil dynamics.

Beyond arousal level, Experiment 1 revealed a phase-dependent effect of slow pupil fluctuations. In the T1-attend condition that produced an auditory attentional blink, probe detection was selectively enhanced during the PD phase occurring 0–2 seconds after a constriction event in slow pupil dynamics below 0.2 Hz. Importantly, Experiment 2 further showed that this phase-dependent enhancement of target detection remained significant after controlling for baseline pupil size, as confirmed by the linear mixed-effects model. These results suggest that the PD phase indexes a transient brain state that may reflect cortical gain modulation across sensory systems, creating a temporal window in which perceptual integration becomes more efficient.

Consistent with this interpretation, the PD phase has been associated with decreased activity in sensory-related regions and increased activity within the default mode network (Yellin et al., 2015). More recent work further suggests that neural stimulus representations themselves may vary as a function of pupil-linked fluctuations (Yang et al., 2026). Notably, this temporal window corresponds closely to the timescale of slow pupil fluctuations linked to large-scale cortical dynamics (Raut et al., 2021, 2025). One possible physiological contributor to such slow fluctuations is vascular oscillators, which generate spontaneous low-frequency changes in arteriolar diameter and can propagate as slow hemodynamic waves (Broggini et al., 2024). Extending these observations beyond the visual domain, the present results demonstrate that ongoing brain-state dynamics indexed by pupil phase can modulate performance in auditory detection tasks. Importantly, these findings indicate that the vulnerability revealed by the attentional blink is not fixed but is continuously modulated by slow transitions in brain state.

Classical models of the attentional blink, including the two-stage model (Chun and Potter, 1995) and the Episodic Simultaneous Type, Serial Token (eSTST) model (Wyble et al., 2009), propose that T1 processing temporarily occupies resources required for working memory consolidation, thereby suppressing the tokenization of a subsequent T2 target. The present findings suggest that part of the trial-to-trial variability (i.e., the probability of successful tokenization) in this process may depend on ongoing brain state fluctuations unfolding over several seconds. The enhanced T2 detection observed during the PD phase of slow pupil fluctuations suggests that the gain of the attentional gating mechanism may be dynamically modulated by global neural state. Consequently, the severity of the attentional blink may depend not only on the temporal structure of the stimulus stream but also on slow fluctuations in neuromodulatory state. From this perspective, attentional capacity may be better understood as an effective capacity that varies dynamically with neuromodulatory activity, potentially linked to LC–NA function. To directly test this hypothesis, Experiment 2 implemented real-time pupil monitoring to trigger stimulus presentation during specific phases of ongoing pupil dynamics. This approach allowed us to monitor the brain’s internal state in real time and manipulate the timing of target presentation while keeping external stimulus parameters constant. Consistent with the observations from Experiment 1, targets presented during the PD phase were detected more accurately than those presented during the PC phase. Although the magnitude of the behavioral improvement was modest, these results provide causal evidence that the phase of slow pupil-linked brain state fluctuations can influence the outcome of temporal attention tasks.

More broadly, this study demonstrates the potential of real-time pupillometry as a method for probing the functional role of ongoing brain state dynamics in cognition, extending recent approaches that detect pupil-phase events during cognitive tasks (Kronemer et al., 2024). Our findings are also consistent with a previous real-time pupil-triggering study demonstrating that fluctuations in pupil dynamics track variations in attentional state (Keene et al., 2022). By aligning stimulus presentation with internal physiological fluctuations, it becomes possible to causally investigate how slow arousal and neuromodulatory dynamics shape cognitive processing. Taken together, our findings suggest that attentional capacity should be understood from a dynamic perspective. The attentional blink emerges not solely from stimulus-driven processing limitations, but from the interaction between rapid sensory processing and ongoing fluctuations in brain state unfolding over several seconds. Future advances in real-time detection and closed-loop stimulus presentation methods may further reveal how slow brain-state dynamics shape moment-to-moment cognitive capacity.

## Acknowledgements

This work was supported by Grants-in-Aid for Scientific Research from the Japan Society for the Promotion of Science (grant number 24K16883).

## Author contributions statement

YS and HL conceived the experiments. YS conducted the experiment(s), analyzed the data, and wrote the manuscript. HL provided feedback and comments on the manuscript. All authors designed the experiments, discussed the analyses, results, and interpretation, and revised the paper.

